# Predictive abstract task representations support few-shot learning in the rat brain

**DOI:** 10.64898/2026.07.26.740301

**Authors:** Mousa Karayanni, Maciej M. Jankowski, Yonatan Loewenstein, Israel Nelken

## Abstract

Few-shot learning reveals core mechanisms of flexible cognition and adaptive decision-making in animal behavior by showing how animals rapidly infer rules, categories, or action strategies from sparse experience. Few-shot learning often depends on abstract representations that capture task structure and generalize across similar instances of the same task. The orbitofrontal cortex (OFC) is known to encode abstract representations, but whether these representations can adapt to new contingencies at the rapid timescale of few-shot learning is unknown. We trained rats on a sequence-learning task with recurring structure but changing contingencies. Rats demonstrated few-shot learning, reaching near-optimal performance within 2-3 rewards after each contingency change, exploiting the task’s recurring structure to rapidly generalize across sequence blocks. We recorded OFC activity using Neuropixels probes during task performance. The firing rate of OFC units in the few seconds preceding each poke encoded the task state and predicted the selected action. Remarkably, this predictive activity also encoded the abstract notion of ‘role’: the function of an action under the current contingencies. Role representations in the OFC changed on the same rapid timescale as the behavioral changes. We suggest that few-shot learning in this task is supported by the rapid remapping of roles onto actions.

## Introduction

Learning is often a gradual process, with improvements in performance requiring many trials and reinforcements (Meister, 2022). In some cases, however, experimental subjects show rapid gains in performance, approaching near-optimal levels after only a few trials. This rapid learning is commonly referred to as few-shot learning. Few-shot learning often relies on generalizing prior knowledge to novel situations (Song et al., 2022; Wang et al., 2021). Through prior experience, concepts that capture structure shared across observations are formed. Novel observations are represented in terms of these concepts, rather than in terms of their low-level sensory properties. The encoding of an observation by such concepts is referred to as an “abstract representation,” and is believed to be a critical component of few-shot learning (Johnston & Fusi, 2023; Song et al., 2022; Wang et al., 2021).

Abstract representations of task structure have been reported in several brain regions (Bernardi et al., 2020; Courellis et al., 2024; El-Gaby et al., 2024; Nieh et al., 2021; Vaidya et al., 2021; Wilson et al., 2014; Zhou et al., 2021). Importantly, such representations have been identified in tasks in which learned rules were generalized to novel examples (Bernardi et al., 2020; Courellis et al., 2024; El-Gaby et al., 2024; Vaidya et al., 2021; Zhou et al., 2021). The OFC is known to be crucial for behavioral flexibility in tasks with frequently changing contingencies (Kim & Ragozzino, 2005). Prior studies have demonstrated that OFC can support abstract representations, but the way these representations develop and change over time has been studied on the slow timescale of sessions and days, over which learning gradually developed. Whether abstract representations in the OFC can be updated on the fast timescale of few-shot learning, reflecting rapidly changing task contingencies, has not been tested.

In the present study, we close this gap. We recorded from OFC in freely moving rats performing a task that required rapid adjustment to changing contingencies. Rats exhibited few-shot learning by leveraging knowledge of task structure. OFC responses encoded abstract representations that flexibly and rapidly changed to reflect current task contingencies at the behavioral time scale.

### Behavior

Experiments were conducted in the Rat Interactive Foraging Facility (RIFF) (Jankowski et al., 2023). The RIFF (Fig. 1a) is a circular arena (diameter: 160 cm) with six interaction areas (IAs, Fig. 1b) around its circumference, each containing two speakers, a water port, and a food port. A ceiling-mounted camera tracked the location of the rat in real time. The RIFF interacted with the behaving animal by sensing its location and actions (movements and pokes into the ports) in real time (30 Hz) and responding with auditory and visual signals, as well as delivering rewards from its IAs.

**Figure 1.**
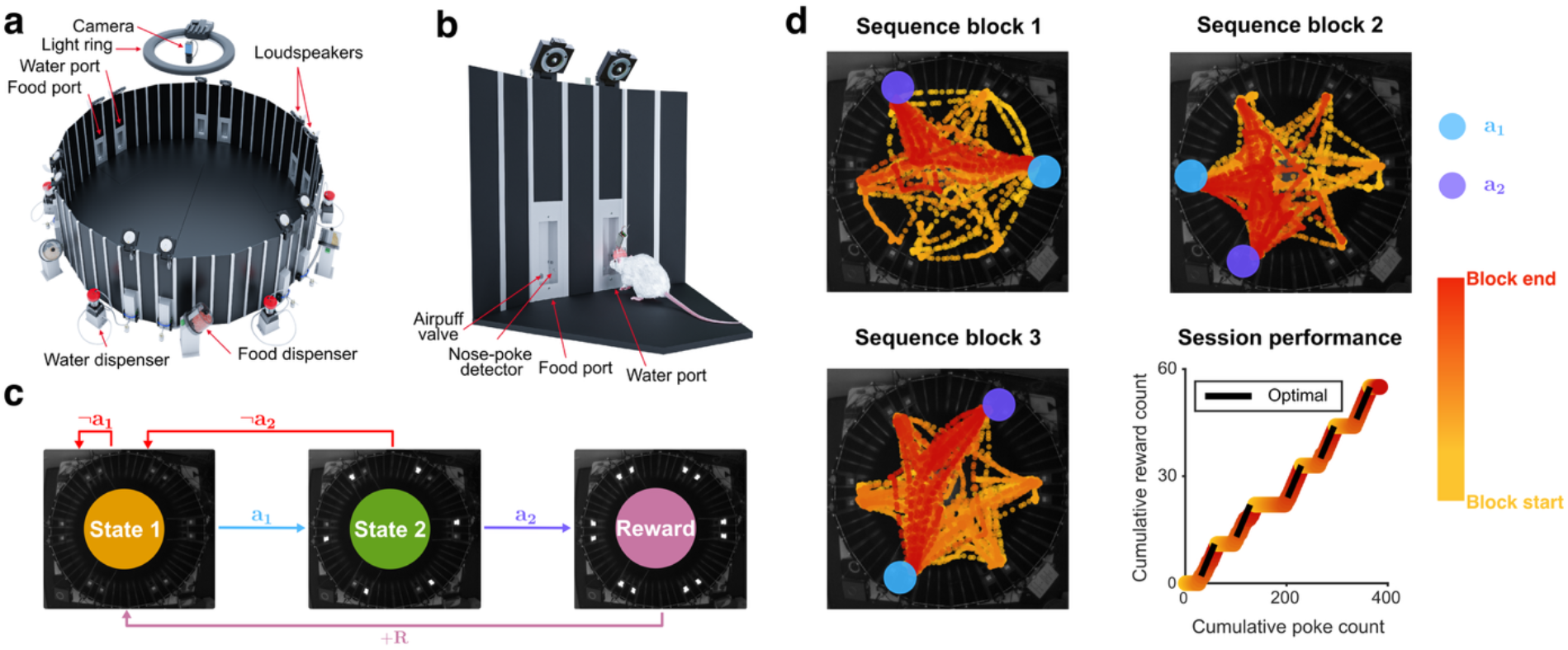
Experimental environment, task, and example behavior. (a) Overview of the Rat Interactive Foraging Facility (RIFF). The 1.6 m open-space arena has six interaction areas (IAs) along its circumference, each with food and water ports and two loudspeakers. Rat location in the arena is identified online using an overhead video camera. (b) Detailed view of an IA, with the rat drawn to scale. (c) Simplified MDP of the sequence-learning task with three states and six actions. *a*_1_ is the correct action in State 1, and *a*_2_ is the correct action in State 2; the other five actions are denoted ∼*a*_1_ and ∼*a*_2_. (d) Behavioral data from an example session. The top two panels and bottom-left panel show behavior in the first three sequence blocks of the session. The rat’s trajectory encodes time using a color gradient from yellow (start of the sequence block) to red (end of the sequence block). The IA of the first correct action (*a*_1_) and of the second correct action (*a*_2_) are marked in cyan and purple, respectively. Early in each block, trajectories reach multiple IAs. Later trajectories visit mostly the two correct IAs, with scattered visits to additional ones, likely used to obtain reward. Bottom right: cumulative performance across the session (five completed sequence blocks). After the second reward in each sequence block (start of black lines), performance is close to optimal (three pokes per reward, slope of the black line).

Rats were trained to sequentially poke in two selected IAs. If successful, the next poke in any of the IAs yielded a reward. Once the rats obtained 11 rewards, the sequence of IAs to poke was changed, without a cue, and the rats had to relearn it. We will refer to the set of pokes associated with a particular sequence as a “sequence block”. Daily sessions included multiple sequence blocks (N=484 sessions with at least one completed sequence block, performed by 5 rats; median: 5 sequence blocks per session; interquartile range: 4-6; see Supplementary Table 1).

Within each sequence block, the task was implemented as a Markov Decision Process (MDP) (Sutton & Barto, 1998). A simplified version of the MDP is shown in Fig. 1c (the MDP is described in its full details in Methods; see Supplementary Fig. 1a). The simplified MDP had 3 states: State 1, State 2, and the reward state, each state marked by distinct auditory and visual cues (see Methods). In each state, the rats could perform one of 6 possible actions, where an action consisted of moving to the center region of the RIFF, followed by poking in one of the IAs. State 1 was the initial state. When in State 1, choosing action *a*_1_ (poking in the first IA of the sequence) resulted in a state transition to State 2. Similarly, choosing action *a*_2_ (poking in the second IA of the sequence) in State 2 resulted in a state transition to the reward state. Any other action in these two states resulted in a return to the initial state, State 1. In the reward state, all actions yielded a reward, followed by a transition to State 1. We will refer to *a*_1_ in State 1 and *a*_2_ in State 2 as the correct actions.

The IA locations of *a*_1_ and *a*_2_ were assigned randomly for each sequence block under two constraints: First, *a*_1_ and *a*_2_ had to be different (so that there were 6 × 5 = 30 distinct sequences in total). Second, two subsequent sequence blocks had to be different (although they could share one of *a*_1_ and *a*_2_). Thus, the first sequence block of a session was randomly chosen from 30 possible sequences, and all subsequent sequence blocks were randomly chosen from 29 possible sequences.

Figure 1d depicts the behavior of one rat in three successive sequence blocks from one session. Early in each block, the rats visited multiple IAs (the yellow part of the trajectory; color indicates time within the sequence block), whereas later their behavior became more stereotyped, with rats visiting in order the two IAs associated with *a*_1_ and *a*_2_, and potentially a third port they used for the reward (the red part of the trajectory). The cumulative rewards for this session are shown in the lower-right panel of Figure 1d (this session continued with two additional completed sequence blocks, for a total of five). At the beginning of each sequence block, as the rat searched for the correct ports, rewards were not being obtained, so the cumulative reward count remained constant (‘exploration’). Later in the block, the rats identified the correct actions and performed them repeatedly in the correct order, resulting in a steady increase in cumulative rewards (‘exploitation’) until the transition to the next sequence restarted exploration.

This behavioral change from exploration to exploitation was abrupt and occurred around the second reward in each sequence block. In the lower-right panel of Figure 1d, the black lines begin at the second reward of each sequence block and have a slope of one reward for every three pokes, the highest possible reward rate. Cumulative reward increased at a similar rate from the second reward onward, suggesting that the rats were already close to optimal performance. This is quantified in Figure 2a, which shows the average number of pokes required to obtain each reward within a sequence block. The first reward required, on average, more than 20 pokes, but the second reward required fewer than 7 pokes on average, and only about 5 pokes were required on average to obtain the third reward. This was already close to the minimum of 3 pokes for a reward. Thus, the rats identified *a*_1_ and *a*_2_ after two successful attempts, demonstrating few-shot learning.

**Figure 2.**
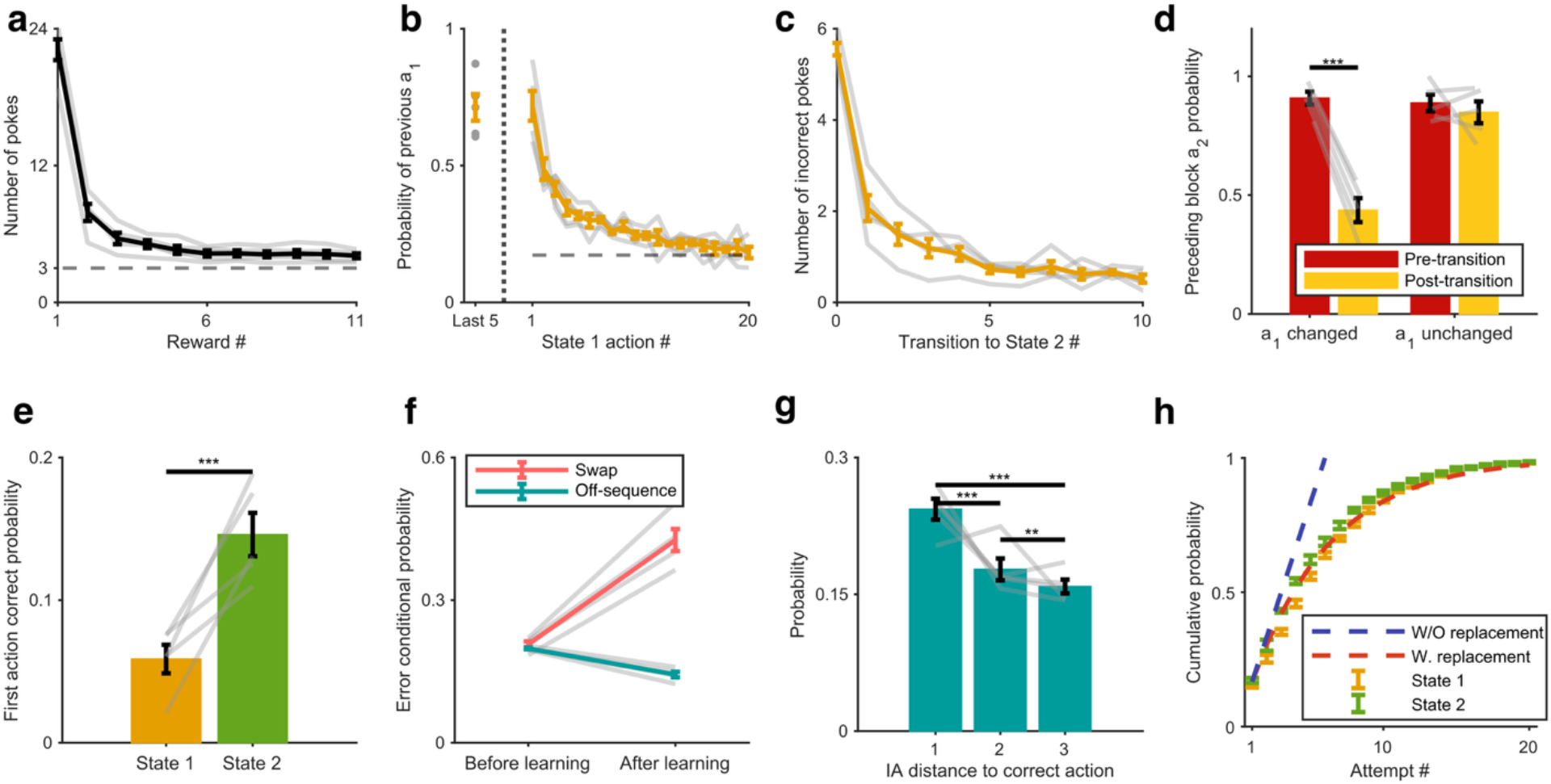
Rats rapidly learned the task using knowledge of the task structure. In (a–g), gray lines show individual rats. (a) Number of pokes per reward for consecutive rewards within a sequence block. The bold line shows the mean across rats, and the error bars denote SEM across rats. Rats approached near-optimal performance (dashed line, 3 pokes per reward) by the third reward. (b) Probability of performing the previous *a*_1_ in State 1 at a sequence-block change. The first point is the average probability over the last five pokes before the sequence-block change. Same convention as (a). The dashed line denotes the probability that *a*_1_ is shared across consecutive sequence blocks; this baseline deviates slightly from 1/6 because of the constraint that the new sequence must differ from the previous one, and is therefore 5/29. (c) Number of incorrect pokes in State 1 as a function of the number of transitions to State 2 before any reward was obtained (that is, before rats identified *a*_2_ of the new sequence block). Same convention as (a). (d) Probability to repeat the previous *a*_2_ in State 2 immediately before and after a sequence-block change, plotted separately for cases in which *a*_1_ changed or remained the same. Analysis is restricted to blocks in which rats identified the new *a*_1_ within three pokes. Mean ± SEM across rats. (e) Probability that the first action in each state was correct, at sequence-block transitions in which both *a*_1_ and *a*_2_ changed. Mean ± SEM across rats. (f) Conditional probability, given an error, that it was a swap error or a poke at a single off-sequence IA (off-sequence probability divided by four), before and after the second reward. Mean ± SEM across rats. (g) Distribution of off-sequence errors after learning as a function of IA distance from the current-state correct action. For completed sequence blocks, we calculated the probability for each distance within each session, averaged across sessions within each rat, and show the mean ± SEM across rats. Error bars show mean ± SEM across rats. (h) Cumulative probability of identifying the correct action in State 1 and State 2 as a function of the number of attempts, shown together with theoretical predictions for sampling with replacement (geometric distribution) and sampling without replacement (linear increase reaching certainty after six attempts). Mean ± SEM across rats.

In the following paragraphs, we characterize learning in this task. We show that (1) rats adapted rapidly to sequence-block changes, (2) they learned the correct actions in the absence of reward (using state transitions instead), (3) they leveraged task structure to generalize across blocks, (4) post-learning errors reflected learned task knowledge, and (5) their search strategy was overall consistent with random sampling but included idiosyncratic search preferences.

### Rats adapted rapidly to sequence-block changes

We examined the probability of poking the previous first IA of the sequence (“previous *a*_1_”; Fig. 2b) immediately after a sequence change. Right after the change, the probability to poke in the previous *a*_1_ was as high as during the last 5 actions of the previous block (Fig. 2b). This indicates that the animals were not aware of the fact that the sequence changed (thus, for example, they likely didn’t count the number of rewards in the block). Importantly, immediately after the first failure to transition to State 2 by performing the previous *a*_1_, the probability of selecting that action dropped towards baseline (Fig. 2b).

### Correct actions were learned from state transitions, even in the absence of reward

To test whether learning required reward, we examined trials in which a correct State 1 poke advanced the rat to State 2, but the following State 2 poke was incorrect, so that no reward was delivered. If learning is contingent on reward, such trials should not change behavior, because no reward was obtained. If, instead, the transition from State 1 to State 2 is itself informative, the probability of performing *a*_1_ on the next visit to State 1 should increase. On average, after a new block began, rats made about 5 incorrect pokes before performing *a*_1_ in State 1, causing a transition to State 2 (Fig. 2c). Note that the next poke in State 2 was incorrect by design in this analysis, and therefore the MDP returned to State 1 without providing the rat with a reward. Critically, upon returning to State 1, the rat performed *a*_1_ again after less than 2 incorrect pokes on average, despite the absence of a reward. Thus, state transitions were sufficient for learning the correct actions.

### Rats leveraged task structure to generalize across sequence blocks

At a sequence block transition, rats could generalize the observation of a change in *a*_1_ to anticipate a change in *a*_2_. When *a*_1_ of the new sequence was different from the previous *a*_1_, the change of the sequence could be detected in State 1, because selecting the previous *a*_1_ failed to cause a transition to State 2. However, in a minority of cases, the new *a*_1_ was the same as the previous *a*_1_, and only *a*_2_ changed in the new sequence. In such cases, rats couldn’t detect the change in the sequence in State 1.

We compared the probability of performing the previous *a*_2_ in State 2 in these two cases: when *a*_1_ had changed (and therefore the rats had the information that *a*_2_ could be different as well) and when it remained the same (and therefore the rats didn’t have information about a sequence change when they got to State 2). In each case, we compared the first State 2 poke of the new block to the State 2 poke that followed the penultimate reward of the previous block. The interpretation of this analysis is complicated by the fact that when *a*_1_ changed, there could be a large number of pokes while the rat searched for the correct *a*_1_ action, potentially causing the rats to forget the previous *a*_2_. We therefore restricted the analysis to blocks in which rats identified the new *a*_1_ within three pokes. The two cases differed significantly (mixed-effects logistic regression with transition, *a*_1_ change, and their interaction as fixed effects, and random intercepts for rat and session-within-rat; interaction p<0.001). As expected, when *a*_1_ remained the same, the previous *a*_2_ was selected as often after the transition as before it (0.87, 95% CI [0.81, 0.92] before vs 0.9, 95% CI [0.83, 0.94] after, p=0.50; Fig. 2d, right). However, when *a*_1_ changed in the new sequence block, the probability of performing the previous *a*_2_ was reduced after the transition (0.91, 95% CI [0.87, 0.94] before vs 0.43, 95% CI [0.34, 0.52] after, p<0.001; Fig. 2d, left).

A direct consequence of this result is that, at sequence transitions where both *a*_1_ and *a*_2_ change, the first attempt in State 1 was overwhelmingly wrong (because it was strongly biased towards the previous *a*_1_, as described in the previous paragraph) while the first attempt in State 2 was less biased towards the previous *a*_2_. Indeed, the probability of performing the correct action on the first attempt was significantly lower in State 1 than in State 2 (only sequence transitions where both *a*_1_ and *a*_2_ changed; mixed-effects logistic regression with rat and session-within-rat as random effects, odds ratio = 3.30, 95% CI [2.38, 4.58], p<0.001; Fig. 2e).

### Errors after learning reflected task knowledge

Because of the abrupt behavioral change that occurred around the second reward, we analyzed errors separately before and after the second reward in each sequence block (’before learning’ and ‘after learning’ below). We distinguished two error types: swap errors, performing the correct action of the other state (*a*_2_ in State 1 or *a*_1_ in State 2; Alleman et al., 2024), and off-sequence errors, pokes that were not part of the correct sequence. We computed both probabilities per session and compared them with a linear mixed-effects model (error type, learning phase, and their interaction as fixed effects, with random intercepts for rat and session-within-rat). The difference between the two error types depended on the learning phase (interaction β = −0.27, 95% CI [−0.29, −0.26], p < 0.001). Before learning, the two probabilities did not differ (β = −0.005, 95% CI [−0.02, 0.01], p = 0.39). After learning, the swap error was significantly more probable than an off-sequence error (β = −0.278, 95% CI [−0.29, −0.27], p < 0.001; Fig. 2f). Moreover, after learning, off-sequence errors were also structured: they tended to occur at IAs that were physically closer to the correct action in each state (linear mixed-effects model for probability as a function of distance with random intercepts for rat and session-within-rat: F(2,1446)=55.4, p<0.001; contrasts: IA distance 1 vs 2, t(1446)=7.54, p<0.001; 2 vs 3, t(1446)=2.60, p=0.009; 1 vs 3, t(1446)=10.1, p<0.001, all passing Benjamini-Hochberg correction at FDR=0.05 Fig. 2g).

### Search strategy consistent with random sampling, with idiosyncratic search preferences

To characterize how rats searched for the correct action in each state, we analyzed pokes made before the correct action was identified and asked whether they were consistent with sampling actions with or without replacement. Under sampling without replacement, the cumulative probability of discovering the correct action as a function of the number of actions is expected to increase linearly and reach certainty after six pokes; under sampling with replacement, it is expected to saturate exponentially. Rat behavior was most consistent with sampling with replacement (Fig. 2h).

Sampling with replacement is usually conceived as selecting the next port randomly from the full set of available options. However, it can also be effectively achieved by rats using a fixed sequence of actions, repeating it during each sequence block as they search for the correct actions. We tested whether rats used preferred sequences of actions during search, examining the first two actions in the first block of each session (restricted to blocks in which the first poke was incorrect, 402 blocks in total). Such sequences could be defined allocentrically (by the identity of the visited ports), or egocentrically, defined by the selected IA relative to the rat’s orientation prior to the poke: for example, a rat might consistently choose the IA to its right, followed by the IA to its left (see Methods). For each candidate action pair, we asked whether it occurred more often than expected under uniform random choice among six ports (one-sided binomial test, Benjamini-Hochberg corrected across the 6² = 36 possible two-action sequences; expected probability per sequence: 1/36). Only one rat showed a significantly preferred allocentric sequence, but each rat had at least one preferred egocentric sequence (see Methods), accounting for 13–40% of search paths across rats. These results suggest that rats exhibit idiosyncratic preferences in their search behavior (Lebovich et al., 2019).

### Neural responses during task performance

Neural activity was recorded using Neuropixels electrodes from three freely-moving, untethered rats performing the task (see Methods; probe placement in Supplementary Fig. 3). Units in the OFC showed clear modulation of activity around poke times (Fig. 3a). We focused on the activity preceding each poke, since this is the window relevant to the decision to poke. Before all subsequent neural analyses, we regressed out motor-related variance from the neural responses and quantified pre-poke activity using a 2-second window preceding each poke (hereafter, *normalized activity* and *pre-poke activity*; see Methods).

**Figure 3.**
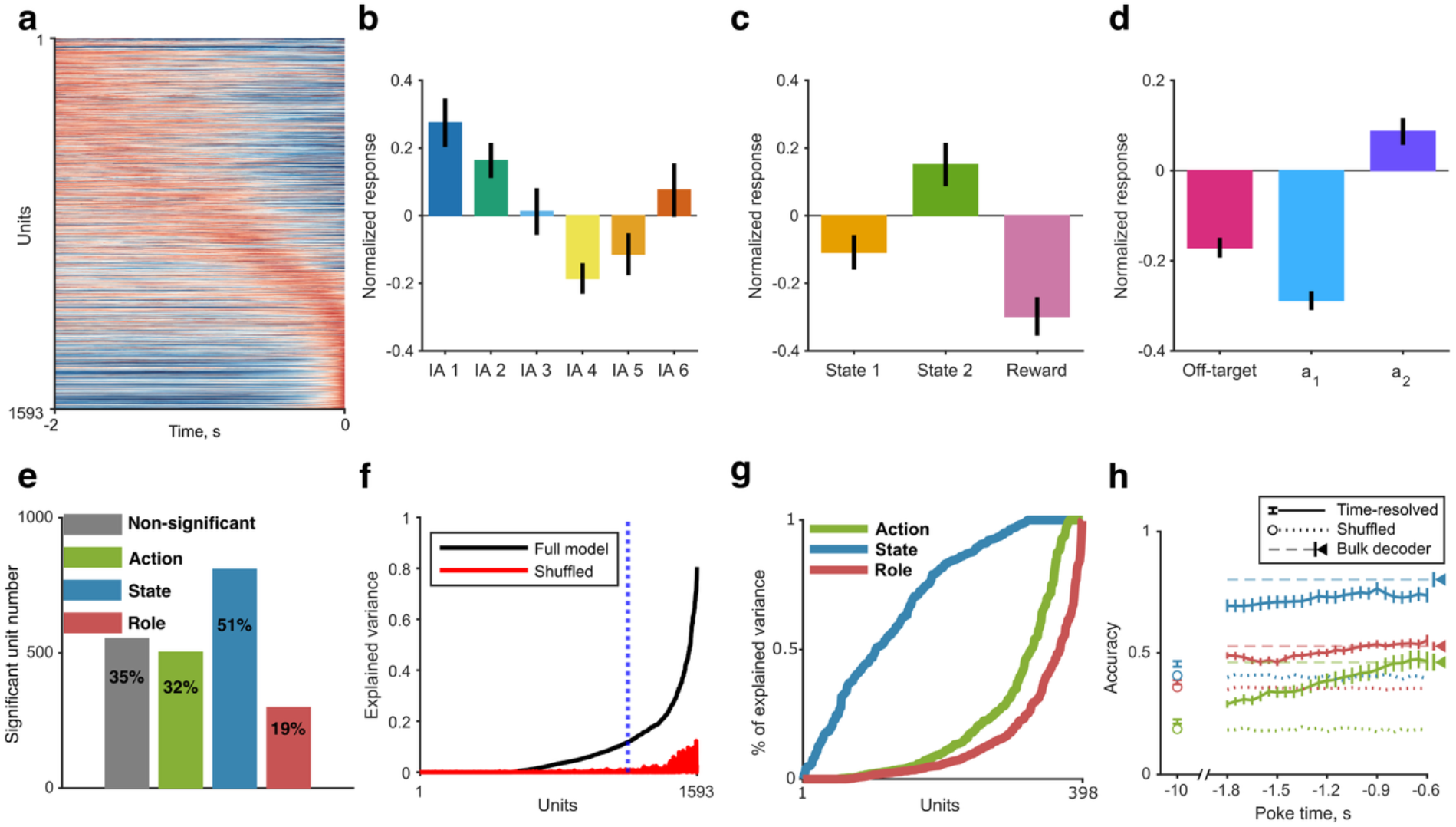
Task-related representations in the OFC population. (a) Average responses of all recorded OFC units around the time of a poke. Each unit’s response was smoothed with a 300 ms moving average and normalized by its peak. Units are ordered by the time of peak activity in the odd-numbered pokes of each session; the activity shown is the average over the even-numbered pokes. (b) The activity of an action-selective unit. Mean ± SEM of the activity preceding pokes for each IA. (c–d) Same as (b), for state-selective and role-selective units, respectively. (e) Counts and proportions of units with significant sensitivity to action, state, and role (units can be sensitive to more than one variable). (f) Explained variance of the full linear model with action, state, and role predictors. Units are sorted by explained variance of the model (black line). The red line shows the explained variance of a model with all predictors shuffled, using the same unit ordering. Blue dashed line marks the top quartile of units by explained variance. (g) Unique explained variance for the top quartile units, shown separately for each predictor; units are sorted by unique explained variance for the corresponding predictor. (h) Decoding accuracy for action, state, and role across pre-poke time windows. Solid lines show time-resolved decoding (mean ± SEM across sessions). Dashed lines show bulk decoder performance (mean ± SEM across sessions; SEM shown at the rightmost point). Dotted lines are at random performance for each variable.

We looked for representations of the states and actions of our MDP, since such representations are often found in OFC (see Methods for details). Figure 3b shows the average pre-poke activity of an action-selective unit for each action. Figure 3c shows a corresponding example of a state-selective unit.

We also asked whether the OFC represents the abstract notion of the *role* of an action within the current sequence-block contingencies. The rats performed a task in which an action could have three roles: in State 1, an action was either the first correct action (*a*_1_) or off-target; in State 2, an action was either the second correct action (*a*_2_) or off-target. In both cases, off-target refers to any action that was incorrect in the current state. We assumed that the representation of the off-target role was shared across the two states. Notably, this role representation was independent of the physical identity of the action: different ports could have the same role in successive sequence blocks. For example, IA 1 might serve as *a*_1_ in one sequence block, whereas IA 4 fulfills that role in the next. If neural activity reflects an abstract role representation, when in State 1, we would expect similar responses to IA 1 in the first block and IA 4 in the second, despite the many physical differences between them. Conversely, the same IA (e.g., IA 1) should elicit different responses across sequence blocks if its role changes (in the example above, its role is *a*_1_ in State 1 in the first sequence block, and therefore it is off-target in State 2 of the same block; it is also off-target in State 1 of the next block). Importantly, if a role representation exists, its rapid association with different actions across sequence blocks could provide a mechanism for few-shot learning. Figure 3d shows an example of a role-selective unit.

Figure 3e summarizes counts and proportions of significant task-related representations among OFC units. About 65% of the units represented at least one task-related variable (action, state, or role). Among the units that showed significant task-related activity, ∼45% exhibited mixed selectivity (selectivity for more than one variable) (Rigotti et al., 2013). This may have happened because the three task-related variables were correlated. Therefore, to capture the unique contribution of each task-related variable to neural activity, we first fitted, for each unit, a linear model predicting average pre-poke activity from all three (main effects only). In Fig. 3f, the black line shows the fraction of variance explained on held-out data using leave-one-out cross-validation, whereas the red line shows the explained variance on the same held-out data for a model in which all predictors were shuffled. As a validation, units that were significantly selective to at least one task-related variable had substantially higher explained variance than units that showed no significant modulation by any task-related variable (‘non-significant units’ in Fig. 3e; Supplementary Fig. 2a). The unique contribution of a variable was defined as the fraction of the full model’s explained variance exclusively attributable to that variable, independently of its correlations with the others (see Methods). We restricted this analysis to units in the top quartile of full-model explained variance (Fig. 3f, blue dashed line). Each variable accounted for positive unique variance in a subset of units (Fig. 3g).

The representation of role could be a consequence of the representation of the value (expected future reward) of an action in a given state. Neural activity in OFC is known to correlate with value (Gottfried et al., 2003; Miller et al., 2022; Padoa-Schioppa & Assad, 2006; Thorpe et al., 1983). Importantly, actions with different roles also have different values. Indeed, when future rewards are temporally discounted, then *a*_2_ in State 2 has a higher value than *a*_1_ in State 1, because *a*_2_ leads directly to a reward, whereas *a*_1_ leads only to State 2 and requires further actions to get a reward. Off-target actions have the lowest value, because they delay future rewards. Thus, value-coding units may show role representation.

To address this critique, we remark that while value is ordinal, role is categorical. Units that code role through their sensitivity to value are expected to have responses that are monotonically ordered, with the activity associated with *a*_1_ intermediate between that for *a*_2_ and off-target. By contrast, categorical role coding does not require such a monotonic ordering, allowing arbitrary response ordering across the three roles. If any ordering of the responses to the three roles is equally probable, only one third of role-representing units would be expected to show the monotonic ordering predicted by value coding.

We restricted the analysis of this prediction to units in the top quartile of the full-model explained variance that also showed positive unique explained variance for role, to ensure that responses to the three role categories were well-estimated. We then tested whether responses to off-target, *a*_1_, and *a*_2_ pokes followed the monotonic ordering predicted by value coding (see Methods). Most of these units (195/336, 58%) showed non-monotonic response profiles (Supplementary Fig. 2b). Thus, the majority of the role-encoding units do not seem to encode value.

Substantial information about task-related variables could be decoded from the OFC population. Decoders trained on the average activity in the 2-second pre-poke window (’bulk decoder’) decoded all three above chance (N = 15 sessions; computed using a shuffled control; probability of correct identification, mean ± SEM; action: 0.46 ± 0.04 vs. 0.18 ± 0.01 for chance, t(14) = 8.13, p < 0.001; state: 0.80 ± 0.03 vs. 0.40 ± 0.01, t(14) = 10.76, p < 0.001; role: 0.56 ± 0.02 vs. 0.36 ± 0.01, t(14) = 11.08, p < 0.001; paired t-tests; Fig. 3h). Information about the task-related variables was present throughout the pre-poke period: all three could be decoded above chance using the activity in 400 ms windows as early as 2 seconds before poke onset and throughout the pre-poke window (see Methods; Fig. 3h).

Using the decoders, we tested our assumption that off-target pokes evoked similar response patterns in States 1 and 2. To do so, we trained a bulk decoder on all pokes in State 1 and then evaluated off-target decoding accuracy on pokes in State 2, relative to chance level (again using a shuffled control). Off-target decoding accuracy exceeded chance level (observed: 0.72 ± 0.03 vs. chance: 0.39 ± 0.06; paired t-test, t(14) = 5.49, p < 0.001; Supplementary Fig. 2d), indicating that the off-target representation was indeed shared across the two states.

### Fast updates of role representations in the OFC

If the OFC contributes to the observed few-shot learning by representing role, then the neurons that represent role should change their responses to actions at sequence transitions. If these changes in neural activity are as rapid as the changes in behavior, they may support few-shot learning.

To track changes in neural activity during learning, we used the finding that rats learned from state transitions (Fig. 2c): in State 1, learning occurred from transitions to State 2, whereas in State 2, it occurred from transitions to the reward state and subsequent reward delivery. In each state, we defined 11 critical pokes whose outcomes could drive learning. In State 1, the first critical poke was the first correct *a*_1_ action after the sequence transition. The subsequent critical pokes in State 1 were defined as the first correct *a*_1_ action after the (*n*−1)-th reward. In State 2, the critical pokes were defined as the correct *a*_2_ action performed just before the *n*-th reward. In the following, we examine the neural activity preceding these pokes. Note that before the first critical poke in either state, the rat did not know that the selected action was correct; this information became available only from the ensuing transition from State 1 to State 2, and from the transition from State 2 to the reward state (and the following reward delivery).

Consider, for example, the role-selective unit shown in Fig. 3d, whose firing was minimal for *a*_1_ in State 1 and maximal for *a*_2_ in State 2. In most cases, when a new sequence block was introduced, the *a*_1_ and *a*_2_ actions of the new sequence were off-target in the previous sequence, and were therefore encoded by an intermediate firing rate in this unit. As learning progressed, we expected responses associated with *a*_1_ in State 1 to decrease from the off-target firing rate to the *a*_1_ firing rate, and the responses associated with *a*_2_ in State 2 to increase from the off-target firing rate to the *a*_2_ firing rate. Indeed, this is what we observed. The cyan line in Fig. 4a shows the pre-poke activity of this unit for the critical pokes of State 1 as learning progressed. Before the first reward, when the animal did not yet know that it was performing *a*_1_ in State 1, the response was at the level of an off-target action (red arrow and dashed line). With subsequent rewards, the activity decreased to the lower values associated with *a*_1_ in State 1 (cyan arrow and dashed line). The purple line shows the corresponding activity for the State 2 critical pokes. Here, too, the response initially resembled that of an off-target action and then increased to the high values associated with *a*_2_ in State 2 (purple arrow and dashed line). In both cases, the change occurred within only 2–3 critical pokes, on a timescale similar to the few-shot learning illustrated in Fig. 2a.

**Figure 4.**
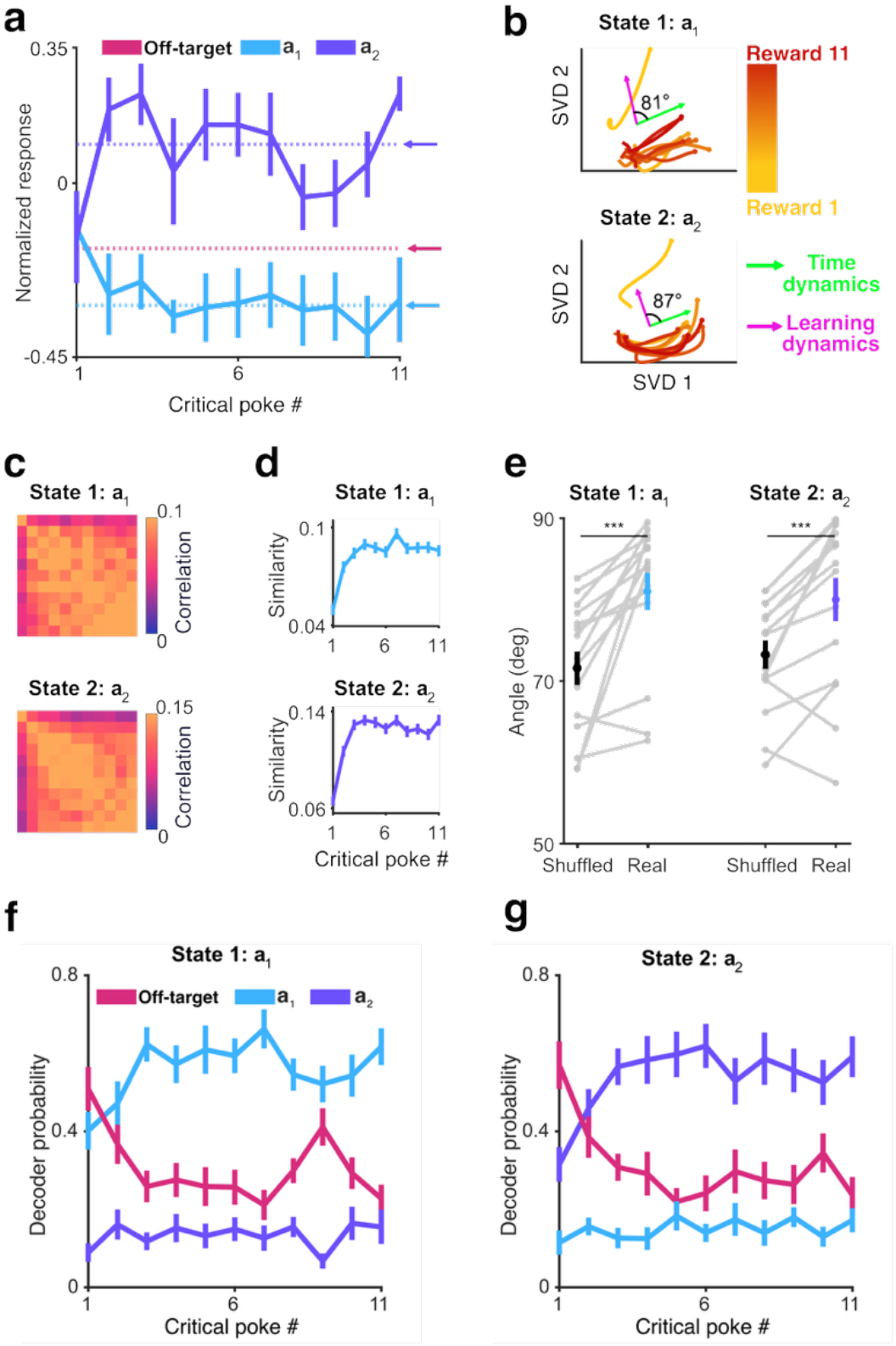
Rapid adaptations in OFC activity track few-shot learning. (a) Rapid changes of role representation in the example unit from Fig. 3d. Responses to *a*_1_ in State 1 and *a*_2_ in State 2 as a function of reward count within a sequence block (averaged across sequence blocks). Arrows and dashed lines indicate the session-averaged response to off-target pokes that occurred after the beginning of a sequence block but before the second reward, and the session-averaged responses to *a*_1_ and *a*_2_ pokes after the second reward. (b) Population trajectories from the same session as the example unit in Fig. 3d and panel a. The population vectors derived from the 2 seconds preceding the same pokes as in (a) are shown projected onto the first two SVD components. The green arrow shows the time-dependent direction; the magenta arrow shows the learning-dependent direction. (c) Average pairwise similarities between population trajectories across sessions. (d) Within-state trajectory similarity as a function of reward count, quantified by the average similarity between each reward-specific trajectory and trajectories from all other reward counts (mean ± SEM across sessions). (e) Acute angle between the learning-dependent and time-dependent directions in State 1 and State 2, compared to a permutation null (mean ± SEM across sessions). Gray lines show individual session values. (f) Bulk role decoder class probabilities for *a*_1_ pokes in State 1 as a function of reward count (mean ± SEM across sequence blocks). (g) Same as (f), for *a*_2_ pokes in State 2.

To assess whether this rapid adaptation also occurred at the population level, we analyzed learning-dependent changes in OFC population trajectories preceding the critical pokes defined above. For each poke, the population trajectory was defined as the time-varying normalized activity of all simultaneously recorded units in the pre-poke window (see Methods). Figure 4b shows two-dimensional projections (computed separately for State 1 and State 2) of these trajectories for the session containing the example unit in Fig. 4a. In both states, the trajectory preceding the first critical poke was clearly different from the subsequent trajectories, which became progressively more similar as rewards accumulated.

To quantify this effect, we computed pairwise similarities between all trajectories (in the original neural space, without dimensional reduction), yielding an 11×11 similarity matrix per sequence block and state. These similarity matrices encode how the population’s neural activity changed as learning progressed within a sequence block, separately for each action in its state. We then averaged these matrices across blocks (Fig. 4c). In both states, the first trajectory was, on average, less similar to the later ones than the later trajectories were to each other. Figure 4d summarizes this effect by showing, for each reward position, the mean similarity to the other ten trajectories. Trajectory similarity rose between the first and second critical pokes in both states (State 1: t(78) = 4.47, p < 0.001; State 2: t(78) = 4.42, p < 0.001). In State 2, similarity also rose between the second and third critical pokes (t(78) = 2.87, p = 0.003). These increases passed Benjamini-Hochberg correction across the ten consecutive comparisons at FDR = 0.05. No later comparison passed correction in either state, consistent with the transition to exploitation after the second reward (Fig. 2a).

This effect could simply reflect slow drifts in neural activity over time. We therefore repeated the analysis on reward pokes, which followed the correct pokes in State 1 and State 2 and were therefore separated by comparable or longer time gaps. Neural trajectories before reward pokes were more similar to each other than State 1 and State 2 trajectories, even at the beginning of a sequence block: the first-to-third similarity increase was significantly smaller for these reward-state pokes than for correct pokes in State 1 and State 2 (0.024 ± 0.008 vs. 0.049 ± 0.005; two-sample t-test, t(235) = 2.75, p = 0.006; Supplementary Fig. 2e), arguing against slow time drift as the explanation.

Interestingly, the learning-dependent changes in the population trajectories were nearly orthogonal to the within-trial time-dependent dynamics (Fig. 4b, 4e; see Methods). For each session, we projected the population trajectories to 3 dimensions using SVD. We estimated the direction in which learning shifted the trajectories by training a linear classifier to separate trajectories preceding the first 2 critical pokes from those preceding critical pokes 3 to 11. We estimated the time-dependent direction of the dynamics by computing the tangent of the population trajectory at each time point and averaging across time points and pokes. We then compared the observed angle between the learning and time-dependent directions to the mean of a null distribution generated for each session by shuffling the critical-poke label of each trajectory, recomputing the learning direction from the shuffled labels. The observed angles were significantly closer to 90° than the shuffled control in both states (Fig. 4e, State 1: 81.0 ± 2.3° vs. 70.3 ± 2.3°, paired t test, t(14) = 4.46, p < 0.001; State 2: 80.0 ± 2.7° vs. 73.2 ± 1.9°, paired t test, t(14) = 3.91, p = 0.0016).

Importantly, the rapid learning-related changes in population activity reflected the emergence of selectivity to role. We used the decoder trained to identify the role of the poke (Fig. 3h) to predict the classes of each of the 11 critical pokes in each state. The neural activity preceding the first critical poke tended to be classified as off-target (red line in Figs. 4f, 4g). However, already at the second critical poke, the classifier’s probabilities increased for the appropriate role (cyan line in Fig. 4f, purple line in Fig. 4g) and decreased for the off-target role, paralleling the rapid learning of the correct actions (Fig. 2a) again.

## Discussion

Rats engaged in a task requiring the repeated learning of action sequences within sessions. They demonstrated few-shot learning -rapid adaptation to new contingencies - exploiting the structure of the task to generalize across sequence blocks.

Remarkably, not all experiences contributed equally to learning. Rats immediately adjusted their behavior after a failed attempt at the previously correct action following a sequence change. Yet failures during exploration did not have the same effect: the cumulative probability of finding the correct action was consistent with search with replacement (even though rats had preferred action sequences; Fig. 2h). We hypothesize that experiences drive learning when their outcomes violate expectations. During exploration, most pokes are expected to fail, and the informative event is the unexpected state transition that occurs when the correct action is selected. In contrast, during exploitation, the selected action is expected to succeed, and the informative event is its unexpected failure. Testing this will require further work.

We studied how neural activity in the OFC encoded three task-related variables: action, state, and role. OFC representations of the first two, action and state, are well-established. Action representations correspond to the representation of future navigational goals reported by Basu et al. (2021). State representations correspond to locations within an abstract cognitive map of the task, which is also represented in the OFC (Moneta et al., 2024; Schuck et al., 2018; Wilson et al., 2014; Zhou et al., 2021). Admittedly, in our task, the state representations could be the consequences, at least in part, of sensory representations: each state was associated with distinct auditory and visual stimuli, and OFC neurons are known to respond to sensory stimuli of both modalities (Thorpe et al., 1983; Winkowski et al., 2018).

The third variable, which we term “role,” is novel. Role captures the function of an action in the current state. It is an abstract representation independent of the action’s physical identity: the same port serves different roles in different sequence blocks. In our task, actions, states, and the state-transition map all remained fixed in all sequence blocks; only the assignment of roles to actions changed. Remarkably, we found that, at sequence block transitions, the OFC population activity encoding role adapted on the same time scale as the behavioral few-shot learning.

The concept of role extends prior work on abstract task representations in OFC. Zhou et al. (2021) showed that when rats learned multiple odor sequences in which the task structure was preserved but the sensory cues changed, OFC eventually encoded the ordinal position of the cue in the sequence rather than its sensory identity, demonstrating that OFC can abstract over sensory input to represent invariant task structure. These abstract representations emerged over multiple sessions. Here we show, in a similar vein, that OFC can abstract over specific actions to represent their function within the stable state-level contingencies. Importantly, we further show that the abstract representation adapts rapidly to novel contingencies.

Value is the variable through which the OFC has been shown to support incremental learning, accumulated over multiple actions (Hattori et al., 2023). Role and value are partially confounded, so we explicitly demonstrated that the majority of the units that showed role sensitivity did not encode value (Supplementary Fig. 2b). Nevertheless, we do not consider value and role as independent variables. Rather, we suggest that value representations may underlie the emergence of role representations.

Indeed, behavioral results show that learners start by forming detailed evaluations of options, but then transition to categorical representations that capture the recurring structure of a problem (Ashby & Maddox, 2005; Chi et al., 1981; Collins & Frank, 2013; Dickinson, 1985; Fitts & Posner, 1967). This is observed, for example, in perceptual learning and face recognition: with experience, subjects perceive stimuli as integrated wholes rather than as collections of features (Gauthier et al., 2000; Tanaka & Farah, 1993). At the neural level, learning is associated with shifts from detailed representations of individual experiences into compressed, abstract representations (Bein & Niv, 2025; Zhou et al., 2021). The resulting representations are more efficient: behaviorally relevant distinctions are preserved while irrelevant details are discarded (Tang et al., 2019).

We interpret the relationship between value and role representations through this lens: role is a categorical representation that groups the behaviorally relevant distinctions while discarding details. We hypothesize that the rats, having performed the task for many months, developed role representation, supporting few-shot learning by reassigning role rather than by incremental updating of the value of each action based on the history of reward accumulation. We therefore predict that recordings from the OFC during early training will reveal value-encoding neurons but fewer (or no) role-encoding neurons, and that role-encoding neurons will emerge later during training.

The emergence of the abstract role representation may occur through a number of computational schemes. The information bottleneck (IB) principle posits optimal representations that preserve information about behaviorally relevant variables while being as simple as possible (Tishby et al., 2000; Tishby & Zaslavsky, 2015). We expect that applying the IB principle to the MDP describing the behavioral task (as in Niediek et al., 2024) may result in the formation of role representations. Alternatively, Johnston & Fusi (2023) offer a computational account for the emergence of abstract representations: training on multiple tasks with shared structure naturally leads to their emergence, in both supervised and reinforcement learning.

Altogether, these results link a behavioral capacity to a likely neural substrate. Rats showed few-shot learning, adapting to new contingencies within a few trials by exploiting the task’s recurring structure. These behavioral manifestations were matched by neural plasticity in the OFC - the representation of role that reassigned actions to their current function - on the same rapid timescale. Role is abstract, grouping actions by their function, independent of their sensorimotor properties. Its rapid reassignment may be the mechanism by which the fixed task structure is used to learn new contingencies from a handful of observations.

### limitations

The present data are correlational. We establish that OFC encodes role and shows changes in role representations on the same time scale as few-shot learning, but whether this representation is necessary for the behavior cannot be determined from recordings alone. Targeted disruption of OFC activity during sequence block transitions would be required to test this. Furthermore, we analyzed the representation of each task-related variable separately. We characterized how action, state, and role are each represented in OFC activity, but the three interact by their nature. Capturing these interactions would require a conceptual framework in which action, state, and role are coupled (Langdon & Engel, 2025). Finally, we recorded extracellular activity and did not identify the cell types or projection targets of the recorded units, so the role representation cannot be assigned to a defined population.

## Methods

### Behavioral task

#### Animals

Five female Sprague Dawley rats (270-325 g) from Envigo LTD, Israel were used for this study. The animals were housed in pairs per cage and had unrestricted access to water and standard rodent food prior to the experiments. The rats were maintained in a temperature and humidity-controlled room on a 12-hour light/dark cycle with lights on from 10:00AM to 10:00PM. The experiments were carried out in compliance with the regulations of the ethics committee at The Hebrew University of Jerusalem. The Hebrew University is accredited by the Association for Assessment and Accreditation of Laboratory Animal Care (AAALAC).

### The rat interactive foraging facility (RIFF)

A full description of the RIFF can be found in Jankowski et al., 2023. In short, the RIFF consists of a large circular arena (160 cm in diameter) with 6 equally spaced interaction areas (IAs), each having a water port, a food port, a pair of LED’s (SPECIAL.090-SE v1.0, LaFayette-Campden Instruments, Loughborough, UK), and two free-field loudspeakers (MF1, Tucker Davis Technologies, Alachua, FL, USA), one above each port. Rat behavior is monitored online using video tracking with a monochrome camera (DMK 33G445 GigE, TheImagingSource, Bremen, Germany) mounted above the center of the arena, with a wide-field lens (T3Z3510CS, Computar, North Carolina, US) to capture the entire arena. The ports were controlled by commercial software (ABET II, LaFayette-Campden Instruments, Loughborough, UK) and a custom-written MATLAB program (MathWorks, Inc., Natick, MA, USA). Each IA contained palatable 45 mg food pellets (F0021, BIO-SERV, Flemington, NJ, USA) in one port and mineral water in the other port. Rats almost invariably poked into the food port.

### Training paradigm and final task

Training consisted of 4 phases that lasted approximately 1.5 – 2 months. During phase 1 (7-10 days), the rats were handled daily by the experimenter. In phase 2 (3-5 days) the rats were habituated to the RIFF for 15-40 minutes every day.

In phase 3, rats were trained to cross an unmarked circle centered on the arena’s midpoint (‘center crossing’) and then to poke into one of the ports, receiving 3 pellets (from a food port) or approximately 0.3 ml of water (from a water port). Rat performance also triggered auditory feedback: a successful center crossing elicited an auditory stimulus which was an excerpt of the word “armed”, frequency-shifted to match the rat auditory system and lasting 200 ms (http://shtooka.net, see Jankowski et al., 2023). This sound was presented from all speakers around the arena. A poke performed immediately following a center crossing elicited a 200 ms, 2 kHz tone from the speaker pair of the IA in which the poke occurred.

During phase 3, the task was progressively made harder. The radius of the central circle decreased daily from 50 cm to 30 cm. The number of rewarding IAs decreased daily from 6 to 1. Once some ports did not provide a reward, the locations of the ports that did changed after a fixed number of rewards. The number of rewards after which the rewarding port location was changed decreased from 41 to 11 rewarded pokes. Phase 3 duration and parameter changes depended on individual rat performance.

Phase 4 incorporated all the states of the final task (Fig. S1a): unarmed State 1 (before center crossing), armed State 1 (following center crossing), unarmed State 2 and armed State 2, and the unarmed reward state and armed reward state. The states were linearly ordered so that the animal could advance from state to state only in the following order: unarmed State 1, armed State 1, unarmed State 2, armed State 2, unarmed reward, armed reward.

Each state was signaled by an auditory and a visual cue. The auditory cue was a frequency-shifted excerpt of a spoken word, presented once every 5 seconds: “cold” in unarmed and armed State 1, “warmer” in unarmed and armed State 2, and “enjoy” in the unarmed and armed reward state. The visual cue was provided by the port LEDs: no LEDs were lit in unarmed or armed State 1; the LEDs on the ports corresponding to the correct IA were lit in unarmed or armed State 2; all LEDs were lit in the unarmed and armed reward state. The transition from an unarmed to an armed state was marked by the “armed” auditory stimulus, presented once after the center crossing.

Rat actions consisted of center crossings and of pokes. To transition from the unarmed to the armed state, rats had to perform a center crossing. A center crossing triggered the same auditory stimulus as in phase 3. Further center crossings performed in an armed state did not elicit any outcome. Pokes in unarmed states did not elicit any outcome. Pokes in an armed state elicited a 200 ms, 2 kHz tone from the two speakers above the poked IA, and caused state transitions. The state auditory cues (i.e., “cold”, “warmer” and “enjoy”) were not played while the rat remained in the center region of the arena.

In armed State 1, one IA was designated correct, and a poke in either port of that IA advanced the animal to unarmed State 2; any other IA poke was incorrect and returned the animal to unarmed State 1. In armed State 2, one IA was designated correct, and a poke in either port of that IA advanced the animal to the unarmed reward state; any other IA poke was incorrect and returned the animal to unarmed State 1. In the reward state, a poke in any port delivered a reward. A correct poke in armed State 2 delivered a reward with a probability that was constant within each session and decreased during training from 75% to 0%. A correct poke in the armed State 1 never resulted in a reward.

The correct action sequence (i.e., the correct IA locations for armed State 1 and armed State 2) changed after a fixed number of rewards that decreased across training from 71 to 11, depending on individual rat performance.

The final task had the same states, transitions, and stimuli as phase 4, with two differences. First, in the final task, in unarmed and armed State 2, the left port LEDs in every IA were lit, regardless of which IA was correct. Thus, the LEDs in State 2 no longer indicated the correct port; instead, they only signaled that the MDP was in State 2. Second, a reward was delivered only following a poke in the armed reward state. The MDP corresponding to the final task is shown in Supplementary Fig. 1a. For simplicity, in the main text we consider an action to consist of a center crossing followed by a poke, causing a transition from one armed state to the next one. We refer to the three state pairs (unarmed and armed) as State 1, State 2, and the reward state. The correct action sequence changed after every 11 rewards.

Rat 1 was the first to be trained on the sequence learning task. Unlike the others, it did not experience any sequence changes during the training phase; these were introduced only after it had progressed to the final task. The sequence changes occurred initially after 71 rewards, and this number was progressively reduced to 11. In the rat’s first 20 sessions with sequence changes, a 200 ms auditory stimulus consisting of a frequency-shifted excerpt of the spoken word “revise” was presented five times in a row. When this rat successfully performed the final task with 41 rewards between sequence changes, it underwent 13 sessions involving a more difficult task, with a sequence that included an additional State 3 (with the same behavior as States 1 and 2, but requiring the identification of a third correct action) before the transition to the reward state. For this rat, only sessions with the standard final task that had length-2 sequences and uncued sequence changes after 11 rewards were included in the analysis.

A circular platform (35 cm radius) was placed at the center of the arena to mitigate the rats’ avoidance of the open space. The platform was covered by a 3D-printed open mesh roof. The platform was present in the arena for all recording sessions. It was introduced during the final phase for rats 1 to 3 and was present from the start of training for rats 4 and 5.

### Surgical procedures

Rats were chronically implanted with Neuropixels 1.0 silicon probes and recorded untethered using the SpikeGadgets Neuropixels headstage. Implantation was performed once each rat had reached stable task performance, 6 months or longer after the start of training.

Animal preparation followed closely the methods of Jankowski et al., 2023. It was performed in two stages: preparation of the implant base, followed by electrode implantation.

To prepare the implant base, the rats were initially anesthetized in an induction chamber with sevoflurane (8% in oxygen, Piramal Critical Care Inc., Bethlehem, PA, USA). The head was shaved, and they were placed in a stereotaxic instrument with a mask for gas anesthesia (David Kopf Instruments, CA, USA). Sevoflurane concentration was slowly adjusted to 2–2.5% and maintained at this level throughout the surgery. A surgical level of anesthesia was verified by the lack of a pedal withdrawal reflex and a slow, stable breathing rate. Body temperature was controlled by a closed-loop heating system with a rectal probe (Homeothermic Monitoring System, Harvard Apparatus, MA, USA). The eyes were protected with sterile saline drops, and the skin on the head was disinfected with a povidone-iodine solution (10%, equivalent to 1% iodine, Rekah Pharm. Ind. Ltd., Holon, Israel). Rats were subcutaneously injected with Meloxicam 5 mg/ml in a dose of about 1.2 mg/kg (Loxicom, Norbrook Laboratories Limited, Newry, Co. Down, Northern Ireland) as well as a local injection of 0.15 ml of Lidocaine at 0.5% W/V (Esracain, Rafa Laboratories ltd, Jerusalem, Israel) to the scalp.

A 1.5–2 cm longitudinal cut of the skin on the head was made. The dorsal surface of the skull was exposed, the connective tissue covering the bones was removed, and the skull bone was treated with a 15% hydrogen peroxide solution (Sigma Aldrich Inc., St. Louis, MO, USA), washed off with sterile saline after approximately 10-20 s. The coordinates for OFC craniotomy site were marked (5-4.2 mm AP, 2 mm ML). Screws (5-7) were placed in the frontal, parietal, and occipital bones. One of these screws was previously soldered with a ground wire and was placed nearest to the craniotomy site.

The screws were first fixed to the bone with resin and then with acrylic dental cement (Super-bond C&B, Sun Medical, Moriyama, Shiga, Japan; Coral-fix, Tel Aviv, Israel), forming the base of the implant. At the selected electrode implantation site, a thin polyimide tube was placed on the skull vertically and cemented with the rest of the implant base. The tube served as a guide to the implantation site for the next surgery. The free end of the ground wire was twisted and covered with a polyethylene cap cemented to the rest of the implant. The wounds were cleaned and treated in situ with antibiotic ointment (synthomycine, chloramfenicol 3%, Rekah Pharm. Ind. Ltd., Holon, Israel). The skin was sutured in the anterior part of the implant with one or two sutures (Nylon, Assut sutures, Corgémont, Switzerland). The skin around the wound was cleaned and covered with a povidone-iodine solution (10%). The rats received an intraperitoneal injection of the antibiotic enrofloxacin (50mg/ml, 5% W/V), at a dose of 15 mg/kg diluted to 1 ml with saline (Baytril, Bayer Animal Health GmbH, Leverkusen, Germany). Meloxicam (Loxicom, Norbrook Laboratories Limited, Newry, Co. Down, Northern Ireland) dissolved in palatable wet food was provided at the home cage for the first 3 days after surgery. The rats were allowed at least 2 weeks of recovery post-surgery before restarting behavioral training.

Prior to the implantation surgery, the electrodes were prepared by soldering a ground wire to the electrode ground and reference contacts. Before surgery, the electrodes and electrode holder were disinfected in a UV sterilizer. The rats were initially anesthetized in an induction chamber with sevoflurane (8% in oxygen). After induction, they were placed in a stereotaxic instrument with a mask for gas anesthesia (David Kopf Instruments, CA, USA). Sevoflurane concentration was slowly adjusted to 2–2.5% and maintained at this level throughout the procedure. Rats received a subcutaneous injection of Meloxicam 5 mg/ml in a dose of about 1.2 mg/kg. The eyes were protected with sterile saline, and body temperature was controlled by a closed-loop heating system with a rectal probe. The dental cement above the implantation site, marked by the polyimide tube, was removed gradually with a dental drill until the skull was exposed. The craniotomy was performed by drilling, and a 0.4-0.8 mm long slit in the dura was gently resected. The electrode was slowly inserted into the brain tissue using a single-axis micromanipulator for a total depth of 5-6 mm (MO-10, Narishige, Tokyo, Japan). The craniotomy was sealed with an elastic silicone polymer (Duragel, Cambridge Neurotech, UK). The ground wire from the electrode and the ground wire from the ground screw were soldered together and covered with acrylic dental cement. Afterward, the electrode was connected to the ZIF connector of the SpikeGadgets Neuropixels headstage, and the headstage assembly was completed. At the end of the surgery, rats received an intraperitoneal injection of enrofloxacin 50mg/ml (15 mg/kg), diluted to 1 ml with saline. Rats were allowed at least 3 days of recovery post-implantation before recordings.

### Histology

Animals were anesthetized in an induction chamber with sevoflurane (8% in oxygen) and then received a lethal injection of sodium pentobarbital (900 mg i.p. per rat, Pentobarbital sodium 200 mg/ml, CTS Chemical Industries Ltd., Kiryat Malachi, Israel). When the rats stopped breathing, they were transcardially perfused in a fume hood with 350 ml of 0.1 M phosphate-buffered saline (PBS, Sigma Aldrich Inc., St. Louis, MO, USA) at room temperature, followed by 400 ml of 7% formaldehyde in 0.1 M PBS at around 4°C (Formaldehyde 35% w/w, Bio-Lab Ltd., Jerusalem, Israel). After perfusion, the brains were removed from the skulls and placed in 7% formaldehyde in 0.1 M PBS for at least 72 h at around 4°C. Following fixation in formaldehyde, the brains were transferred to 30% sucrose in 4% formaldehyde solution in 0.1 M PBS at approximately 4°C (Sigma-Aldrich Inc., St. Louis, MO, USA) for cryoprotection for 7-14 days. The brains were levelled, and a small triangular longitudinal cut on the right hemisphere cortex was made (hemisphere marker). Each brain was placed in a polyethylene embedding mold (Peel-A-Way, Sigma Aldrich Inc., St. Louis, MO, USA). The mold was filled with optimal cutting temperature compound (OCT compound, Tissue-Tek, Alphen aan den Rijn, Netherlands). The polyethylene mold with the brain and OCT was placed in a custom aluminum freezing mold filled with a small amount of absolute ethanol (J.T.Baker, Deventer, Netherlands). The aluminum was then filled with dry ice to ensure that the brain froze rapidly. Frozen blocks with the levelled brains were transferred to a cryostat (CM1950, Leica). Consecutive 35 µm thick coronal sections were cut around the implantation site (left hemisphere). Brain slices were stained with green fluorescent Nissl stain (NeuroTrace® 500/525 Green Fluorescent Nissl Stain, Molecular Probes™, Eugene, OR, USA) according to the manufacturer’s instructions. The slices were then mounted onto slides and covered with a mounting medium (Vectashield H1200, Vector Laboratories, Inc., Burlingame, CA, USA) and a cover glass. The slides were examined under standard fluorescent and bright field microscopy with 4x and 10x objectives.

### Electrophysiology

#### Denoising and spikesorting

Raw electrophysiological data were first denoised with a custom procedure that removed electrical noise shared across channels. A weighted average of the cross-channel activity was subtracted from each channel, followed by a modified version of the DSS algorithm of de Cheveigné & Simon (2008) to remove residual shared noise. The denoised data were spike-sorted with Kilosort 2.5.1 (Pachitariu et al., 2016) and manually curated with Phy2 (https://github.com/cortex-lab/phy). Spike times were binned at 1 ms resolution. Analyses included both single units and multi-units.

### Tagging OFC units

The shank trajectory often spanned multiple histological sections, with each section containing only a short segment of the track. We traced the shank in every section in which it was visible, yielding a set of discrete points. A single representative section was selected as the reference, and the traced points were aligned to it by anatomical landmarks. Because the shank was curved, we fitted these points with a third-order polynomial to reconstruct the full, continuous electrode trajectory. Unit positions were estimated from their relative depth along this reconstructed trajectory. Units were assigned to the OFC using the rat brain atlas of Paxinos & Watson (2007); units outside the OFC were excluded from analysis. Supplementary Table 2 reports the number of recording sessions in each rat, including the total number of units, single units, multi-units, OFC units, and OFC single units included in the analysis.

### Normalized activity and pre-poke activity

Because rats moved freely during the task, we removed movement-related variance from the ongoing OFC activity before all subsequent analyses. The spiking time series of each unit was first smoothed with a 200 ms boxcar filter using zero-phase filtering, and then z-scored across time. The z-scored values were regressed against cos(head direction), sin(head direction), and running speed. We defined the normalized activity as the residual of this model.

Pre-poke activity was derived from the normalized activity. For each poke, we computed the mean normalized activity in the window from 1.8 to 0.2 s before poke onset. Because the activity had been smoothed with a 200 ms zero-phase filter, this measure captured neural activity over the 2 seconds preceding each poke. The window also avoided contamination from the preceding poke, since the minimum inter-poke interval in the sessions used for neural analysis was 2 seconds (Supplementary Fig. 1b).

### Data analysis and statistics

#### Tracking animal posture

During sessions, online camera tracking was used to monitor the rats’ position in the arena by identifying their center of mass (see Jankowski et al., 2023). Post-hoc tracking of additional body landmarks: the base of the implant, neck, and tail, was performed using DeepLabCut v2.2.1 (Mathis et al., 2018), applied to each frame of the RIFF recordings captured at 30 Hz. Rat speed was computed as the temporal derivative of the center-of-mass position, while movement direction was estimated based on the angle formed by the line connecting the base of the implant to the base of the neck.

### Preferred allocentric or egocentric search patterns

For each session, we analyzed the first two pokes of the first sequence block. We restricted this analysis to blocks in which the first poke was incorrect, so that the rat had not yet received any information about the correct action, and the two pokes reflected exploration rather than exploitation. We described each two-poke sequence in two reference frames. In the allocentric frame, a sequence was defined by the identities of the two poked IAs. In the egocentric frame, a sequence was defined relative to the rat’s position at the moment of the center crossing. The ports were referenced to the IA closest to that position, and identified by their distance from it, coded as {−2, −1, 0, 1, 2, 3}. Each pair of pokes, therefore, yielded one allocentric sequence and one egocentric sequence. We counted how many times each observed sequence occurred and asked whether any particular two-poke sequence occurred more often than expected by chance. Under the null hypothesis, the rat selected uniformly at random among the six ports at each step, so that the probability of any specific two-poke sequence was p₀ = 1/6². For a sequence that occurred k times out of n total sequences, we computed a one-sided binomial p-value, P(X ≥ k) with X ∼ Binomial(n, p₀), giving the probability of observing at least k occurrences by chance. Because we tested all 6² possible two-poke sequences, we applied a Benjamini-Hochberg correction (for each rat and reference frame separately) to control the false discovery rate at 0.05.

### Behavioral analyses and statistics

Behavioral comparisons used mixed-effects models with rat and session-within-rat as random effects. Binary outcomes (Fig. 2d, 2e) were modeled with logistic mixed-effects regression fit on individual pokes scored as 1 or 0. For Fig. 2d, we report the probability for each combination of transition and *a*_1_ change with its 95% confidence interval, obtained by transforming the corresponding linear combination of fixed-effect estimates and the endpoints of its confidence interval from the log-odds scale with the logistic function. For Fig. 2e, we report the odds ratio and its 95% confidence interval as the exponentiated fixed-effect estimate and confidence interval. Session-level probabilities (Fig. 2f, 2g) were modeled with linear mixed-effects regression. Comparisons with a 2×2 design (Fig. 2d, 2f) were fit as a single model including both factors and their interaction; the simple effects we report are contrasts within that model.

### Statistical test for significance

To quantify the selectivity of each unit, we computed a one-way ANOVA F-value for each of three categorical task variables, using the unit’s pre-poke response on each poke. Action (IA; six levels) and state (three levels) were evaluated over all pokes. For role, we defined three categories: *a*_1_ in State 1, *a*_2_ in State 2, and off-target. Role was evaluated only on State 1 and State 2 pokes.

We assessed the significance of each F-value with a permutation test. For action and state, labels were permuted across all pokes. For role, labels were permuted within State 1 and within State 2 separately, so that the shuffle preserved state identity. A unit was considered significantly selective for a variable if the p-value was below 0.01.

### Explained variance

To quantify how much of the variance in the pre-poke responses of individual units was uniquely attributable to action, state, and role, we fit a linear regression model with categorical coding of state, action, and role. For reward-state pokes, all three role predictors were zero. The model was fit by ridge-regularized ordinary least squares (λ = 10⁻⁶). Before fitting, each unit’s mean response over the training pokes was subtracted, and the coefficients were fit to these mean-subtracted responses. The training mean was added back to obtain predictions for the held-out poke. The explained variance of the model was computed by leave-one-poke-out cross-validation, estimated as 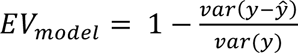, where *y* are the observed responses, *ŷ*. are the predictions on held-out data, and *var* denotes the variance across pokes. As a control, we computed the explained variance of a model in which all three predictors were shuffled across pokes, using the same cross-validation procedure (red line, Fig. 3f). The variance of the held-out prediction errors could be larger than the variance of the responses; in these cases, the explained variance of the model was set to 0.

The unique explained variance of a variable was defined as the reduction in the explained variance of this model when the tested variable was made uninformative. In practice, we compared the full model to a surrogate model in which the predictors related to the tested variable were shuffled, leaving the others intact. For action and state, labels were shuffled across all pokes. For role, labels were shuffled within State 1 and within State 2 separately, leaving reward state pokes unchanged. The explained variance of the surrogate model (*EV_s_*_ℎ*uffled*_) was computed by leave-one-poke-out cross-validation using the same procedure as for the full model. This shuffle was repeated 50 times, and the control explained variance was averaged across repetitions. The unique explained variance of a task-related variable was 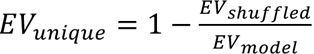. When negative, it was set to 0, and when larger than 1, it was set to 1.

### Role and value

This analysis was restricted to units in the top quartile of full-model cross-validated explained variance that also had positive unique explained variance for role. For each unit, we estimated the rate difference between the activity preceding a correct action and off-target actions. This was performed separately in each state, using ordinary least-squares models and the relevant subset of pokes. The State 1 model used off-target and *a*_1_ pokes; the State 2 model used off-target and *a*_2_ pokes. Each model included action predictors and a single role indicator, set to 1 for the correct action pokes and 0 for off-target pokes. With this encoding, the coefficient of the role indicator estimated the required rate differences, controlled for action. To keep the design matrix full-rank, we used only actions that occurred with at least two pokes at both roles in the corresponding state: off-target and *a*_1_ in State 1, off-target and *a*_2_ in State 2. Because each model was fit within a single state, this contrast was not confounded by state selectivity.

A unit was classified as monotonic if the two contrasts had the same sign and the *a*_2_ contrast was larger in magnitude than the *a*_1_ contrast, that is, sign(*a*_1_ − off-target) = sign(*a*_2_ − off-target) and |*a*_2_ − off-target| > |*a*_1_ − off-target|. This corresponds to the ordering off-target < *a*_1_ < *a*_2_ or off-target > *a*_1_ > *a*_2_. All other units were classified as non-monotonic.

### Decoding task-related variables from neural activity

For the time-resolved decoder, we trained a separate decoder at each pre-poke time point: −10 s, as well as −1.8 s to −0.6 s in 50 ms steps. At each time point, a population vector was formed by summing the normalized activity of each unit over a 400 ms window starting at that time point. The time points from −1.8 s to −0.6 s, the 400 ms summing window, and the 200 ms filtering window were chosen so that the processed data do not depend on neural activity outside the 2 s window preceding each poke. For each time point and target variable, we used leave-one-poke-out cross-validation. Decoding multiple classes used an error-correcting output code (ECOC) classifier with pairwise linear discriminant learners and posterior probability estimation (MATLAB fitcecoc). For the shuffled control, we used surrogate data in which we permuted the training labels of the population vectors, breaking the correspondence between neural activity and labels, repeating the permutation 10 times.

We also performed a bulk decoding analysis, using the pre-poke activity of each unit. Bulk decoding of action, state, and role used the same cross-validation and classifier, with 50 shuffles for the control.

Role decoding, in both the time-resolved and bulk analyses, was restricted to non-reward pokes, since role is defined only in State 1 and State 2.

Because role is defined within state and is therefore correlated with it, role decoding could reflect state information. To control for this, we used a control in which role labels were permuted within state. Role decoding remained above this within-state control for both decoders (bulk decoder: 0.56 ± 0.02 vs. within-state shuffle 0.46 ± 0.01, t(14) = 7.99, p < 0.001; Supplementary Fig. 2c).

### Population trajectory analyses

Population trajectories were constructed from normalized OFC activity. For each of the 11 critical pokes used to track learning of *a*_1_ in State 1 or *a*_2_ in State 2, we extracted the activity of all simultaneously recorded units in the pre-poke window at 1 ms resolution. We refer to the resulting unit-by-time matrix as the population trajectory preceding that poke. We included in this analysis sequence blocks with at least five rewards. To quantify similarity between trajectories, we computed the Pearson correlation of the trajectories for every pair of pokes (separately for each state and each sequence block). These pairwise similarity matrices were then averaged across sequence blocks and sessions, separately for each state.

The low-dimensional visualization (Fig. 4b) and the angle analysis (Fig. 4e) used averaged trajectories. For each state, we averaged the trajectories across sequence blocks, separately for each critical poke. Each unit’s averaged trajectory was then smoothed with a 400 ms boxcar kernel by zero-phase filtering. We then subtracted, for each unit, its mean across the 11 averaged trajectories. We stacked the 11 averaged trajectories of each unit at a given state along the time dimension into a single matrix, with time points across the 11 critical pokes (time points × critical pokes) as rows and units as columns. We applied singular value decomposition to this matrix and projected each of the 11 averaged trajectories onto the first three right singular vectors. Fig. 4b shows the first two dimensions for one example session.

To test whether learning-dependent and time-dependent changes in population activity spanned distinct directions, we defined two vectors in the low-dimensional space for each session and state. The learning axis was obtained by training a linear discriminant classifier (MATLAB fitcdiscr, pseudoLinear) to separate all time points of the early reward trajectories (rewards 1 and 2) from those of the later reward trajectories. The temporal axis was the mean tangent vector, obtained as the mean displacement between consecutive time points, averaged across trajectories and time. We measured the acute angle between the two axes as arccos(|cos θ|), where cos θ is their cosine similarity. To assess significance, we generated for each session a null distribution of 100 angles. Each was obtained by permuting the assignment of critical pokes within each block and repeating the angle estimation. We then compared the observed angle of each session to the mean of its null distribution with a paired t-test across sessions.

---

**Supplementary Figure 1.**
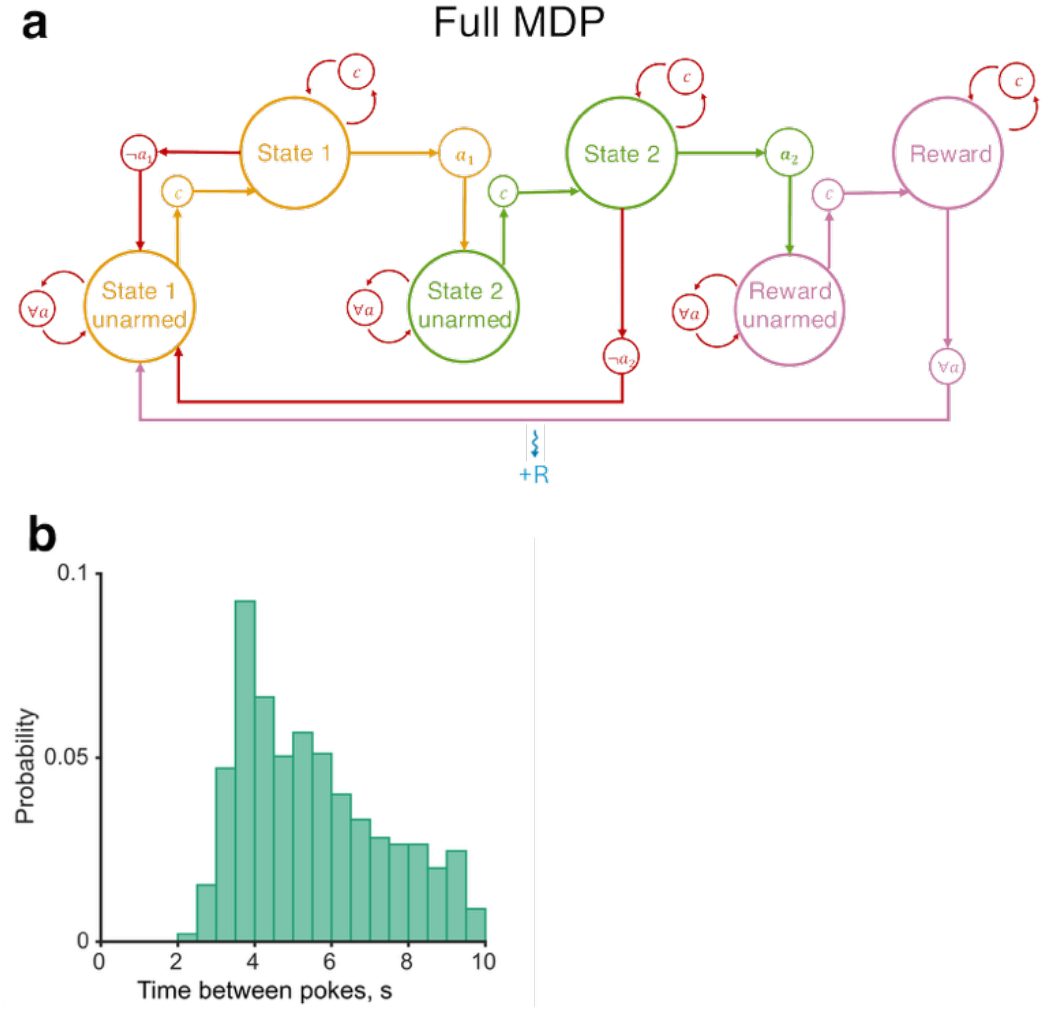
(a) Full MDP of the task, including unarmed states and center-cross actions leading to the armed states. (b) Distribution of inter-poke intervals across all sessions.

**Supplementary Figure 2.**
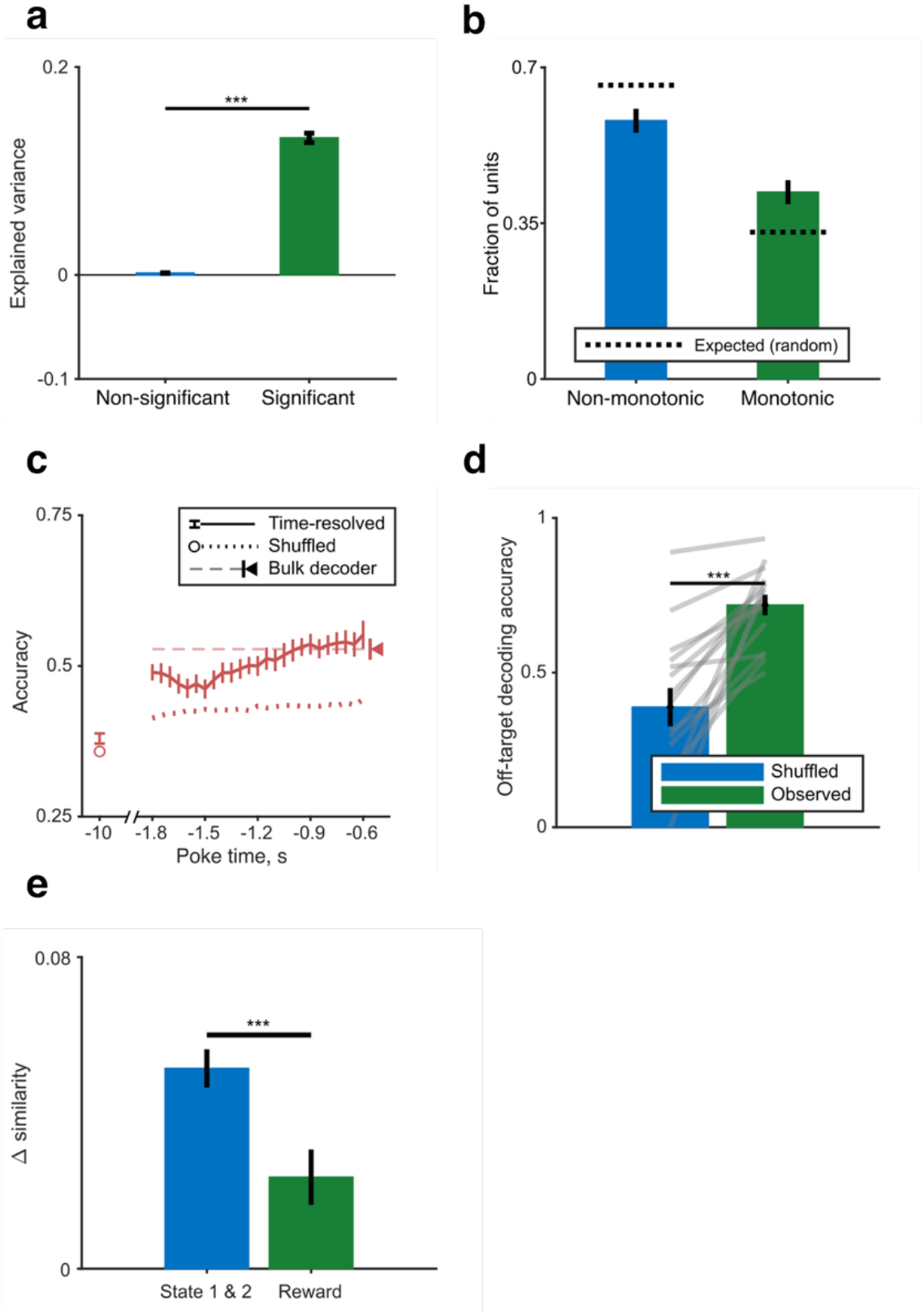
(a) Full-model explained variance for units that were not selective to any task-related variable, compared to units that were significantly selective to at least one variable. (b) Fraction of units with monotonic versus non-monotonic role response profiles (control for the representation of value). Error bars indicate 95% binomial confidence intervals on the proportion. Computed for units in the top quartile of explained variance with positive unique explained variance for role. (c) Role decoder performance as in Fig. 3h, with random performance computed by shuffling role labels only within state. (d) Cross-state generalization of off-target decoding. The role decoder was trained on all State 1 pokes (*a*_1_ vs. off-target) and tested on State 2 pokes (*a*_2_ vs. off-target); gray lines show values for the observed data and random label permutations for individual sessions. (e) Change in population trajectory similarity from the first to the third reward, shown for correct pokes pooled across State 1 and State 2 and for reward state pokes (time-drift control).

**Supplementary Figure 3.**
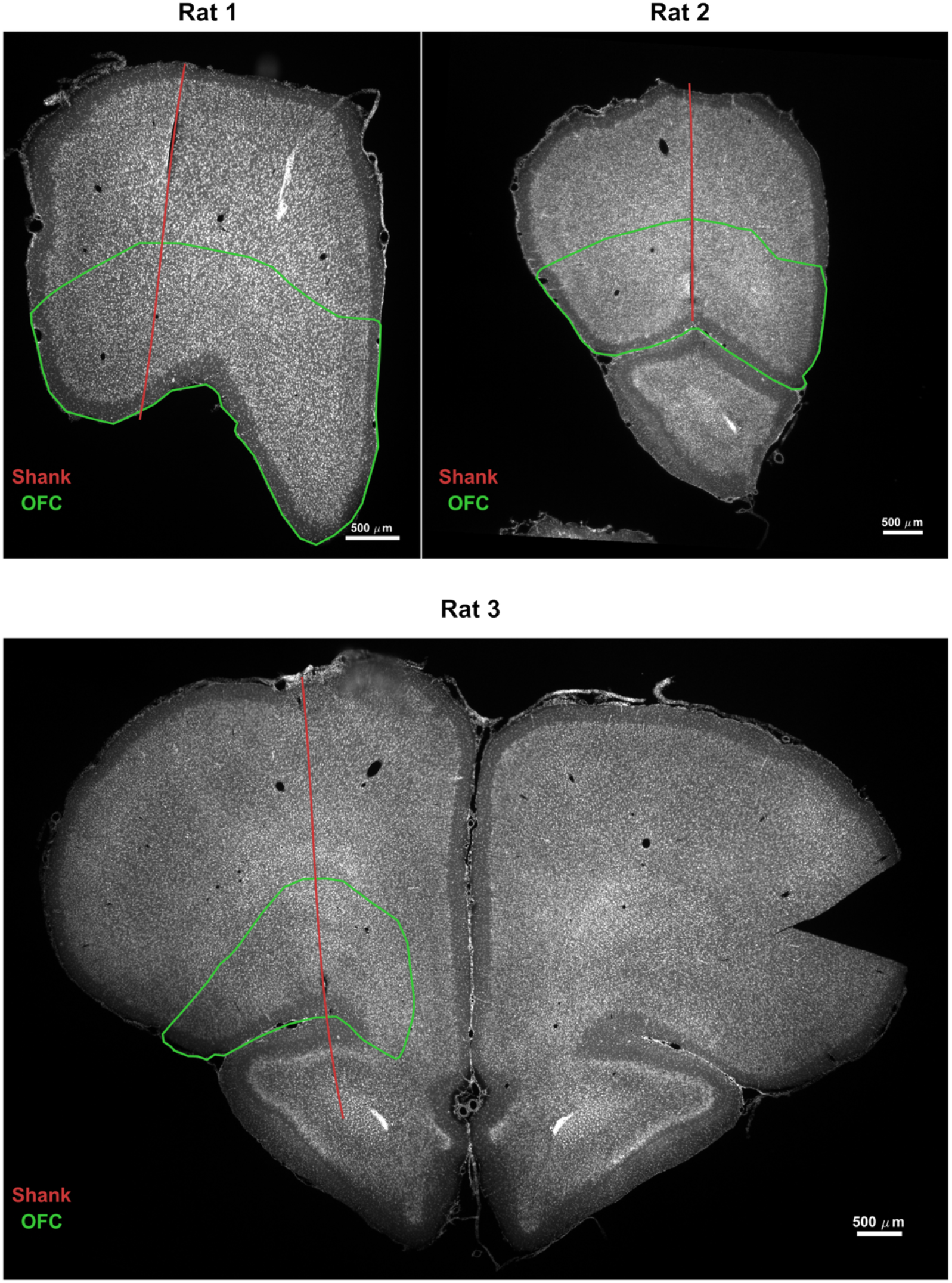
Probe placement in the three recorded rats. Each panel shows a representative coronal section from one rat, with the reconstructed shank trajectory overlaid as a red line and the OFC boundary drawn in green. The shank trajectory was traced across all sections in which it was visible and reconstructed as a continuous curve aligned to the reference section (see Methods). Units were assigned to the OFC using the atlas of Paxinos and Watson (2007).

**Supplementary Table 1.**

| Rat | Total sessions | Total blocks completed | Session duration median [IQR] (min) | Blocks per session median [IQR] |
| --- | --- | --- | --- | --- |
| 1 | 149 | 696 | 94 [87-100] | 5 [3-6] |
| 2 | 63 | 160 | 99 [90-109] | 2 [2-3] |
| 3 | 120 | 436 | 98 [91-113] | 4 [3-5] |
| 4 | 108 | 403 | 108 [95-117] | 4 [3-4] |
| 5 | 44 | 153 | 109 [97-126] | 3 [3-4] |

**Supplementary Table 2.**

| Rat | Session count | Total units | Single units | Multi units | OFC units | OFC single units |
| --- | --- | --- | --- | --- | --- | --- |
| 1 | 6 | 1217 | 528 | 689 | 979 | 432 |
| 2 | 3 | 791 | 235 | 556 | 291 | 81 |
| 3 | 6 | 572 | 165 | 407 | 323 | 94 |

